# Associations between environmental breast cancer risk factors and DNA methylation-based risk-predicting measures

**DOI:** 10.1101/446484

**Authors:** Minyuan Chen, Ee Ming Wong, Tuong L Nguyen, Gillian S Dite, Jennifer Stone, Graham G Giles, Melissa C Southey, John L Hopper, Shuai Li

## Abstract

**Background:** Genome-wide average DNA methylation (GWAM) and epigenetic age acceleration have been suggested to predict breast cancer risk. We aimed to investigate the relationships between these putative risk-predicting measures and environmental breast cancer risk factors.

**Methods:** Using the Illumina HumanMethylation450K assay methylation data, we calculated GWAM and epigenetic age acceleration for 132 female twin pairs and their 215 sisters. Linear regression was used to estimate associations between these risk-predicting measures and multiple breast cancer risk factors. Within-pair analysis was performed for the 132 twin pairs.

**Results:** GWAM was negatively associated with number of live births, and positively with age at first live birth (both P<0.05). Epigenetic age acceleration was positively associated with body mass index (BMI), smoking, alcohol drinking and age at menarche, and negatively with age at first live birth (all P<0.05), and the associations with BMI, alcohol drinking and age at first live birth remained in the within-pair analysis.

**Conclusions:** This exploratory study shows that lifestyle and hormone-related breast cancer risk factors are associated with DNA methylation-based measures that could predict breast cancer risk. The associations of epigenetic age acceleration with BMI, alcohol drinking and age at first live birth are unlikely to be due to familial confounding.

## Introduction

DNA methylation has been investigated as a risk factor for breast cancer. Peripheral blood DNA hypermethylation at the known breast cancer susceptibility genes *BRCA1* and *ATM* has been found to be associated with elevated breast cancer risk [1-3]. Several genome-wide studies of DNA methylation have suggested that peripheral blood DNA methylation, at both global and site-specific levels, is associated with breast cancer risk [4-7].

Two DNA methylation-based measures have been suggested to *could* predict breast cancer risk. One measure is a global DNA methylation measure, genome-wide average DNA methylation (GWAM), which is defined as the average methylation value across probes [8,9]. Two nested case–control studies measured GWAM in pre-diagnosis peripheral blood using the Infinium HumanMethylation 450K BeadChip assay and found that a standard deviation (SD) decrease in GWAM is associated with a 45–65% increased breast cancer risk [4,5]. These results are consistent with global hypomethylation being associated with carcinogenesis, probably by inducing genomic instability [10], and consistent with previous findings that conventional global blood DNA methylation measures, such as *LINE-1* and LUMA, are inversely associated with breast cancer risk [11,12]. A similar measure, the median methylation value across probes, has been found to be prospectively associated with risks of mature B-cell neoplasms [13], urothelial cell carcinoma [14] and prostate cancer [15].

Another measure that has been suggested to *could* predict breast cancer risk is the epigenetic age acceleration, which is the difference between chronological age and biological age based on DNA methylation (DNAm age). While several estimators of DNAm age have been developed [16], the Horvath [17] and Hannum estimators [18] have received the most attention. Recently, a new estimator, known as the PhenoAge estimator, has been developed and uses the ‘phenotypic age’, a linear combination of 10 clinical biomarkers associated with the hazard of aging-related mortality, as a surrogate measure for biological age [19]. Epigenetic age acceleration measures based on both the Horvath and PhenoAge estimators have been found to be associated with incident breast cancer risk: a one-year increase in epigenetic age acceleration measured in pre-diagnosis blood is associated with a 4% increased risk of breast cancer [19,20]. Epigenetic age acceleration has also been found to be associated with risks of other cancers, such as kidney cancer and mature B-cell neoplasms [21].

Both of these putative DNA methylation-based breast cancer risk-predicting measures have environmental, i.e., non-genetic, determinants. We performed a large twin and family study of approximately 2,300 individuals and found that the variance of GWAM across the lifespan is largely determined by environmental factors [8]. To our knowledge, no study has investigated which environmental factors might be associated with GWAM. The variance of epigenetic age acceleration has also been suggested to be influenced by environmental factors [22]. Lifestyle factors, such as smoking, alcohol drinking, obesity, exercise and diet [23,19,24-26], and hormone-related breast cancer risk factors, such as age at menarche and age at menopause [27,28], have been found to be associated with epigenetic age acceleration.

Given evidence above, we hypothesized that conventional environmental breast cancer risk factors are associated with the novel DNA methylation-based risk-predicting measures. We performed an exploratory study to investigate which environmental breast cancer risk factors might be associated with GWAM or epigenetic age acceleration.

## Material and methods

### Study sample

The sample comprised women from the Australian Mammographic Density Twins and Sisters Study [29]. The analytical sample of 479 women, including 66 monozygotic twin (MZ) pairs, 66 dizygotic twin (DZ) pairs and 215 sisters of twins from 130 families, aged 40-78 years and had a mean (SD) age of 55.7 (7.9) years. None of the women sampled had breast cancer.

### GWAM and epigenetic age acceleration calculation

GWAM and epigenetic age acceleration were calculated using DNA methylation data measured by the Infinium HumanMethylation 450K BeadChip assay; see Li *et al.* [22] for more details about the methylation measurement. In brief, DNA was extracted from dried blood spots. Data were normalized using Illumina’s reference factor-based normalization methods and subset-quantile within array normalization [30] for type I and II probe bias correction. An empirical Bayes batch-effects removal method, *ComBat* [31], was applied to minimise technical variation across batches.

GWAM was calculated as the average methylation beta-value across all autosomal probes. We calculated three measures of epigenetic age acceleration based on the DNAm ages estimated by the Horvath, Hannum and PhenoAge estimators, respectively. The Horvath DNAm age was calculated using the online calculator (https://labs.genetics.ucla.edu/horvath/dnamage/), and the Hannum and PhenoAge DNAm ages were calculated as weighted sums of methylation beta-values at 71 probes and 513 probes, respectively, with weights being the reported regression coefficients from the original publications [18,19]. Epigenetic age acceleration was calculated as the residuals from a linear regression model of DNAm age on chronological age and cell counts (CD4T cells, CD8T cells, natural killer cells, B cells, monocytes, granulocytes); therefore, it is independent of chronological age and cell counts. Cell counts were estimated using the Houseman method [32].

### Data collection for breast cancer risk factors

We studied several established and putative breast cancer risk factors, including lifestyle factors and hormone-related factors. All risk factors were collected via telephone-administered questionnaire survey. Lifestyle risk factors included body mass index (BMI), smoking status (non-smoker, former smoker, current smoker), smoking intensity measured as pack-years and alcohol use (non-drinker, former drinker, current drinker). Hormone-related risk factors including age at menarche, parity (nulliparous, parous), number of live births, age at first live birth, oral contraceptive use (non-user, former user, current user), length of oral contraceptive use in years, menopausal status (premenopausal, post-menopausal), age at menopause, hormonal replacement therapy (HRT) use (nonuser, former user, current user) and duration of HRT use in years.

### Statistical analyses

We assessed the correlations between the three epigenetic age acceleration measures using the Pearson’s correlation coefficient. We investigated the associations between GWAM and each epigenetic age acceleration measure using a linear regression model, in which GWAM was the dependent variable and epigenetic age acceleration measure was the independent variable. Each model was adjusted for age, log transformed BMI, smoking status and estimated blood cell counts. To account for the relatedness between family members, the regression model was fitted using the Generalised Estimating Equations (GEE) method with family as cluster.

Associations with breast cancer risk factors were analysed separately for each of the risk-predicting measures (GWAM and the three epigenetic age acceleration measures) using a linear regression model, in which the risk-predicting measure was the dependent variable and the risk factor was the independent variable. The model was fitted using GEE, and adjusted for age, log transformed BMI, smoking status and estimated blood cell counts, with the exception that smoking status was dropped from the model investigating pack-years of smoking, and estimated cell counts were dropped from the model investigating the epigenetic age acceleration measures.

Right-skewed continuous risk factors (BMI, pack-years of smoking, number of live births, and durations of HRT and oral contraceptive use) were log transformed to have approximately normal distributions. The analyses for pack-years of smoking, age at menopause, number of live births, age at first live birth, length of HRT use and length of oral contraceptive use, were restricted to women who were ever smokers, post-menopausal, parous, ever HRT users and ever oral contraceptive users, as appropriate. Given our study was exploratory, no multiple testing adjustment was performed, and a *P-*value of 0.05 was used as the nominal statistical significance level.

We performed within-pair analyses using data from the 132 twins to investigate associations after controlling for the confounding effects of factors (both known and unknown) that are shared by twins. Within-pair differences in risk-predicting measures, risk factors and covariates were calculated, and used to estimate within-pair associations using an ordinary linear regression model without an intercept. The within-pair analyses were performed for all twin pairs, MZ pairs and DZ pairs. We investigated the heterogeneity in within-pair association by zygosity through testing an interaction term of the risk factor with zygosity in the regression model for all twin pairs.

## Results

Table 1 shows the descriptive statistics of the investigated risk factors for the 479 women. For lifestyle risk factors, the median (inter-quartile range [IQR]) BMI was 26.9 (22.9–29.8) kg/m^2^. There were 188 (39.2%) ever smokers with a median (IQR) pack-years of 7.0 (2.3–16.0) years, and 287 (59.9%) ever alcohol drinkers. For hormone-related risk factors, the mean (SD) age at menarche was 13.0 (1.5) years. There were 431 (90.0%) parous women with a mean (SD) age at first live birth of 24.9 (4.9) years and a median (IQR) number of live births of 3.0 (2.0–4.0), 400 (83.5%) oral contraceptive ever users with a median (IQR) use length of 3.6 (0.5–9.0) years, 328 (68.5) post-menopausal women with a mean (SD) age at menopause of 46.1 (6.1) years, and 186 (38.8%) HRT ever users with a median duration of use of 4.0 (1.0–8.1) years.

**Table 1.** Association estimates between DNA methylation-based risk-predicting measures and breast cancer risk factors

| Risk factors | Descriptive statistic | Genome-wide average DNA methylation |  | Horvath epigenetic age acceleration measure |  | Hannum epigenetic age acceleration measure |  | PhenoAge epigenetic age acceleration measure |  |
| --- | --- | --- | --- | --- | --- | --- | --- | --- | --- |
| | | $\beta$ (SE)<sup>a</sup> | *P* | $\beta$ (SE) | *P* | $\beta$ (SE) | *P* | $\beta$ (SE) | *P* |
| Body mass index (kg/m<sup>2</sup>)<sup>b</sup> | 26.9 (22.9–29.8) | –6.06 (6.97) | 0.38 | 2.37 (1.16) | 0.04 | 1.48 (1.15) | 0.20 | 4.69 (1.74) | 0.01 |
| Smoking<sup>c</sup> |  |  |  |  |  |  |  |  |  |
| Non-smoker | 291 (60.8) | Reference |  | Reference |  | Reference |  | Reference |  |
| Former smoker | 147 (30.7) | –3.91 (2.82) | 0.17 | 0.21 (0.52) | 0.68 | –0.26 (0.45) | 0.56 | 0.75 (0.62) | 0.22 |
| Current smoker | 41 (8.6) | –1.98 (4.37) | 0.65 | 0.53 (0.72) | 0.46 | –0.94 (0.59) | 0.11 | 2.04 (1.07) | 0.06 |
| Pack-years of smoking<sup>b</sup> | 7.0 (2.3–16.0) | 0.28 (1.53) | 0.86 | 0.12 (0.24) | 0.61 | 0.21 (0.17) | 0.21 | 0.92 (0.27) | 7.3E-04 |
| Alcohol drinking<sup>c</sup> |  |  |  |  |  |  |  |  |  |
| Non-drinker | 192 (40.1) | Reference |  | Reference |  | Reference |  | Reference |  |
| Former drinker | 52 (10.9) | 2.75 (4.23) | 0.51 | –0.29 (0.83) | 0.73 | –1.15 (0.61) | 0.06 | –0.20 (0.82) | 0.81 |
| Current drinker | 235 (49.1) | –2.19 (2.95) | 0.46 | 1.06 (0.49) | 0.03 | 0.39 (0.42) | 0.36 | 1.25 (0.63) | 0.047 |
| Age at Menarche<sup>d</sup> | 13.0 (1.5) | 0.93 (0.80) | 0.24 | 0.31 (0.15) | 0.04 | 0.13 (0.12) | 0.27 | 0.41 (0.19) | 0.03 |
| Parity<sup>c</sup> |  |  |  |  |  |  |  |  |  |
| Nulliparous | 48 (10.0) | Reference |  | Reference |  | Reference |  | Reference |  |
| Parous | 431 (90.0) | 3.73 (4.04) | 0.36 | –0.58 (0.76) | 0.45 | –0.36 (0.76) | 0.64 | –1.08 (1.11) | 0.33 |
| Number of live births<sup>b</sup> | 3.0 (2.0–4.0) | –7.83 (3.07) | 0.01 | 0.02 (0.64) | 0.97 | 0.28 (0.45) | 0.52 | –0.42 (0.71) | 0.56 |
| Age at first live birth (years)<sup>d</sup> | 24.9 (4.9) | 0.81 (0.33) | 0.01 | –0.07 (0.06) | 0.20 | –0.11 (0.05) | 0.02 | –0.02 (0.07) | 0.72 |
| Oral contraceptive use (years)<sup>c</sup> |  |  |  |  |  |  |  |  |  |
| Non-user | 79 (16.5) | Reference |  | Reference |  | Reference |  | Reference |  |
| Former user | 367 (76.6) | –2.88 (3.32) | 0.39 | 0.17 (0.58) | 0.77 | –0.91 (0.54) | 0.09 | –0.25 (0.75) | 0.74 |
| Current user | 33 (6.9) | –3.02 (5.67) | 0.59 | 0.15 (0.86) | 0.86 | –1.57 (0.92) | 0.09 | 0.30 (0.75) | 0.69 |
| Length of oral contraceptive use (years)<sup>b</sup> | 3.6 (0.5–9.0) | –1.55 (1.01) | 0.12 | –0.04 (0.20) | 0.83 | –0.20 (0.17) | 0.24 | 0.33 (0.36) | 0.36 |
| Menopause<sup>c</sup> |  |  |  |  |  |  |  |  |  |
| Pre-menopausal | 151 (31.5) | Reference |  | Reference |  | Reference |  | Reference |  |
| Post-menopausal | 328 (68.5) | 5.52 (4.10) | 0.18 | –0.49 (0.65) | 0.45 | 0.73 (0.68) | 0.28 | 0.09 (0.77) | 0.91 |
| Age at menopause (years)<sup>d</sup> | 46.1 (6.1) | 0.10 (0.30) | 0.74 | –0.05 (0.05) | 0.29 | –0.03 (0.04) | 0.46 | –0.03 (0.07) | 0.64 |

Table 1 continued
| Risk factors | Descriptive statistic | Genome-wide average DNA methylation |  | Horvath measure of epigenetic age acceleration |  | Hannum measure of epigenetic age acceleration |  | PhenoAge measure of epigenetic age acceleration |  |
| --- | --- | --- | --- | --- | --- | --- | --- | --- | --- |
| | | $\beta$ (SE)<sup>a</sup> | *P* | $\beta$ (SE) | *P* | $\beta$ (SE) | *P* | $\beta$ (SE) | *P* |
| Hormonal replacement therapy use<sup>c</sup> |  |  |  |  |  |  |  |  |  |
| Non-user | 293 (61.2) | Reference |  | Reference |  | Reference |  | Reference |  |
| Former user | 138 (28.8) | 2.78 (3.31) | 0.40 | 0.02 (0.55) | 0.98 | 0.42 (0.50) | 0.40 | −0.25 (0.75) | 0.74 |
| Current user | 48 (10.0) | 4.57 (4.66) | 0.33 | −0.17 (0.71) | 0.81 | 0.77 (0.68) | 0.25 | 0.30 (0.75) | 0.69 |
| Length of Hormonal replacement therapy use (years)<sup>b</sup> | 4.0 (1.0–8.1) | −0.12 (1.53) | 0.94 | 0.22 (0.29) | 0.45 | 0.34 (0.23) | 0.13 | 0.33 (0.36) | 0.36 |

<sup>a</sup> Values are expressed as the original $\beta$ (SE) × 10,000; <sup>b</sup> Values are expressed as median (inter-quartile range); <sup>c</sup> Values are expressed as number of women (percentage); <sup>d</sup> Values are expressed as mean (standard deviation).

The mean (SD) GWAM was 0.53 (0.003). The mean (SD) Horvath, Hannum and PhenoAge measures of epigenetic age acceleration were 0 (4.7), 0 (4.2) and 0 (6.2), respectively. The three epigenetic age acceleration measures were moderately correlated: r = 0.63 between the Horvath and Hannum measures, r = 0.53 between the Horvath and PhenoAge measures, and r = 0.44 between the Hannum and PhenoAge measures (all P<0.05). GWAM was not found to be associated with age (β and 95% confidence interval [CI] are reported as the original β and 95% CI multiplied by 10,000, the same for the GWAM results below; β=0.34, 95% CI: –0.01, 0.69), nor with any epigenetic age acceleration measure (the Horvath measure: β = −0.36, 95% CI: –0.87, 0.14; the Hannum measure: β = –0.20, 95% CI: –0.82, 0.42; the PhenoAge measure: β = –0.27, 95% CI: –0.71, 0.16).

GWAM was found to be associated with number of live births and age at first live birth: women with more live births had a lower GWAM (β = –7.83, 95% CI: –13.85, –1.81), and women with an older age at first live birth had a higher GWAM (β = 0.81, 95% CI: 0.16, 1.45). No associations were found between GWAM and the other risk factors (Table 1).

When assessed using the Horvath epigenetic age acceleration measure, women were an additional 0.24 (95% CI: 0.01, 0.47) years older for each 10% greater in BMI, and current alcohol drinkers were 1.13 (95% CI: 0.26, 1.99) years older than non-or former drinkers. Each additional year in age at menarche was associated with an additional 0.31 (95% CI: 0.01, 0.61) years.

The Hannum epigenetic age acceleration measure was found to be associated with age at first live birth; each additional year in age at first live birth was associated with a 0.11 (95% CI: 0.01, 0.20) years reduction.

When assessed using the PhenoAge epigenetic age acceleration measure, women with 10% greater BMI were an additional 0.47 (0.13, 0.81) years older, women with 10% greater pack-years of smoking were an additional 0.09 (95% CI: 0.04, 0.15) years older, and current alcohol drinkers were 1.30 years older (95% CI: 0.16, 2.44) than non-drinkers or former drinkers. Each additional year in age at menarche was associated with an additional 0.41 (95% CI: 0.04, 0.78) years.

The 10 combinations of risk-predicting measures and associated risk factors found above were included in the within-pair analyses; see Table 2 and Figure 1. For all twin pairs, evidence of a within-pair association was found for the Horvath epigenetic age acceleration measure with BMI (β = 6.44, 95% CI: 1.79, 11.09) and alcohol drinking (current alcohol drinkers compared with non-drinkers or former drinkers: β = 1.72, 95% CI: 0.08, 3.36), and for the Hannum epigenetic age acceleration measure with age at first live birth (β = –0.16, 95% CI: –0.31, –0.02). The Hannum epigenetic age acceleration measure was also found to be associated with age at first live birth in the analysis for MZ pairs only (β = –0.21, 95% CI: –0.38, –0.03). No evidence of a within-pair association heterogeneity between MZ and DZ pairs was found (all P>0.05).

**Figure 1.**
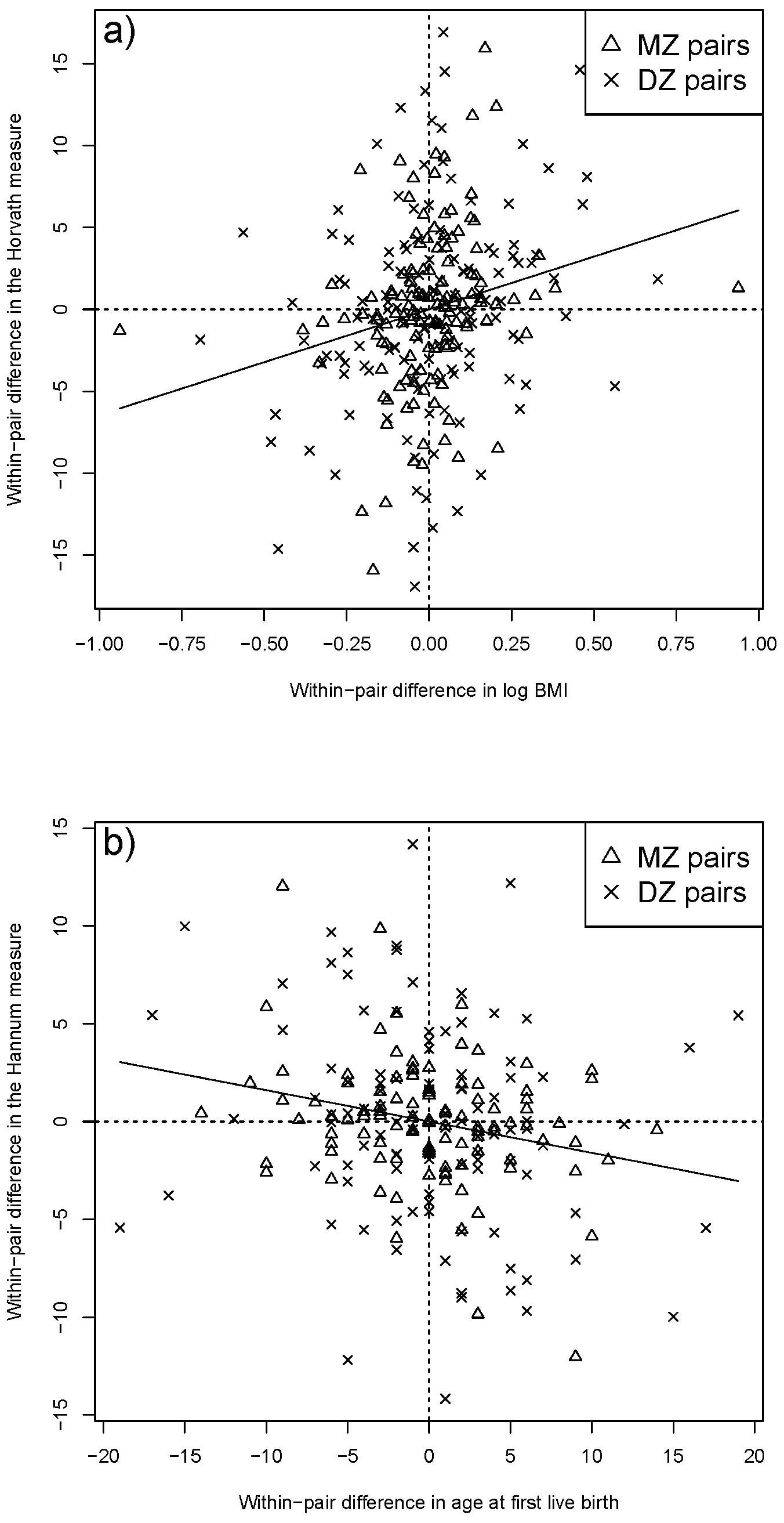
Scatterplots of the observed within-pair associations. a) Within-pair difference in the Horvath epigenetic age acceleration measure vs the difference in BMI. b) Within-pair difference in the Hannum epigenetic age acceleration measure vs the difference in age at first live birth. Solid lines are fitted regression lines. Each twin pair was plotted twice, with both positive and negative within-pair differences.

**Table 2.** Association estimates in within-pair analyses for DNA methylation-based risk-predicting measures and breast cancer risk factors

| Risk-predicting measure – risk factor | All pairs |  | Monozygotic twin pairs |  | Dizygotic twin pairs |  | P for heterogeneity |
| --- | --- | --- | --- | --- | --- | --- | --- |
|  | β (95% CI) | P | β (95% CI) | P | β (95% CI) | P |  |
| Genome-wide average DNA methylation <sup>a</sup> |  |  |  |  |  |  |  |
| Number of live births | 0.73 (−12.92, 14.38) | 0.92 | 1.32 (−19.69, 22.33) | 0.90 | −0.74 (−22.07, 20.58) | 0.95 | 0.77 |
| Age at first live birth | −0.04 (−1.13, 1.04) | 0.94 | 0.12 (−1.82, 2.05) | 0.91 | −0.58 (−2.19, 1.04) | 0.49 | 0.53 |
| Hovarth epigenetic age acceleration measure |  |  |  |  |  |  |  |
| BMI | 6.44 (1.79, 11.09) | 7.5E-03 | 5.78 (−1.02, 12.58) | 0.10 | 6.95 (0.42, 13.48) | 0.04 | 0.87 |
| Current alcohol drinkers vs non- or former drinkers | 1.72 (0.08, 3.36) | 0.04 | 0.37 (−2.08, 2.82) | 0.77 | 2.61 (0.33, 4.88) | 0.03 | 0.29 |
| Age at menarche | 0.22 (−0.42, 0.86) | 0.50 | 0.27 (−0.85, 1.39) | 0.64 | 0.18 (−0.66, 1.02) | 0.67 | 0.72 |
| Hannum epigenetic age acceleration measure |  |  |  |  |  |  |  |
| Age at first live birth | −0.16 (−0.31, −0.02) | 0.03 | −0.21 (−0.38, −0.03) | 0.02 | −0.17 (−0.40, 0.07) | 0.17 | 0.99 |
| PhenoAge epigenetic age acceleration measure |  |  |  |  |  |  |  |
| BMI | 4.55 (−1.06, 10.15) | 0.11 | 4.25 (−3.94, 12.45) | 0.31 | 4.70 (−3.20, 12.60) | 0.25 | 0.97 |
| Pack-years | 0.52 (−0.66, 1.71) | 0.40 | 0.47 (−0.97, 1.91) | 0.53 | 0.58 (−1.69, 2.86) | 0.63 | 0.94 |
| Current alcohol drinkers vs non- or former drinkers | 1.79 (−0.19, 3.77) | 0.08 | 1.39 (−1.55, 4.33) | 0.36 | 2.05 (−0.75, 4.85) | 0.16 | 0.79 |
| Age at menarche | 0.19 (−0.58, 0.96) | 0.63 | 1.32 (0.01, 2.64) | 0.05 | −0.20 (−1.22, 0.82) | 0.70 | 0.13 |

<sup>a</sup> Values are expressed as the original $\beta$ (95% CI) $\times 10,000$ .

## Discussion

In an exploratory study of women aged 40-78 years, we investigated cross-sectional associations between two putative DNA methylation-based breast cancer risk-predicting measures, GWAM and epigenetic age acceleration, and several environmental breast cancer risk factors. We found that GWAM was associated with hormone-related risk factors (number of live births and age at first live birth), while epigenetic age acceleration was associated with lifestyle risk factors (BMI, smoking and alcohol drinking) and hormone-related risk factors (age at menarche and age at first live birth). We also found evidence that the associations of epigenetic age acceleration with BMI, alcohol drinking and age at first live birth remained after controlling for familial factors shared by twins.

Following our previous finding that the variance in GWAM is determined by environmental factors across the lifespan [8], here we found that certain hormone-related risk factors including number of live births and age at first live birth were associated with GWAM. To the best of our knowledge, our study is the first to report that GWAM is associated with these two environmental factors. Paradoxically, we found that lower GWAM, which is potentially associated with an elevated risk of breast cancer [4,5], was associated with more live births and younger age at first live birth, which protect against breast cancer. One implication of this paradox is that parity might not modify breast cancer risk through changing GWAM. Obviously, our analyses are exploratory, our findings require replication by other studies of similar design.

Similar to previous studies [23,19,24-26], we also found epigenetic age acceleration being positively associated with lifestyle risk factors including BMI, smoking and alcohol drinking. These results suggest that an unhealthy lifestyle could accelerate biological ageing, consistent with both epigenetic age acceleration and an unhealthy lifestyle potentially being associated with an increased breast cancer risk. We, thus, hypothesize that an unhealthy lifestyle might modify breast cancer risk through accelerating biological ageing. Further work is required to test this hypothesis.

The PheoAge epigenetic age acceleration measure, compared with the other two epigenetic age acceleration measures, appears to more strongly associated with lifestyle risk factors: the PhenoAge measure was associated with all three studied factors, while the other two measures were associated with fewer. Similar findings have arisen from previous studies; see Table 1 from Horvath *et al.* [33]. These differences in associations might be due to the difference in the development of these DNAm age estimators: the Horvath and Hannum estimators are trained on chronological age [17,18], while the PhenoAge estimator is trained on the ‘phenotypic age’, which is strongly associated with mortality [19].

We found that epigenetic age acceleration was associated with hormone-related breast cancer risk factors, as has been previously found [27,28]. Our study is the first to suggest that age at first live birth is associated with the Hannum epigenetic age acceleration measure. Investigating epigenetic age acceleration and pubertal development, Binder and colleagues found a negative association between the Horvath measure and age at menarche [27], which is in the opposite to the direction of association we observed. One difference between the two studies is that Binder *et al.* studied adolescent girls while we studied older women. Importantly, previous findings about the association between the Horvath epigenetic age acceleration measure at adolescence and children pubertal development are inconsistent: no association was reported by the Avon Longitudinal Study of Parents and Children [34], whereas a positive association was reported by a Finnish study [35]. Additionally, older the PhenoAge epigenetic age acceleration measure was also found to be associated with older age at menarche in our study, providing more evidence in terms of the direction of association between epigenetic age acceleration for older women and age at menarche.

Results from the within-pair analyses suggest that our observed associations of the Horvath epigenetic age acceleration measure with BMI and alcohol drinking, and of the Hannum epigenetic age acceleration measure with age at first live birth are unlikely to be due to confounding of familial factors (both known and unknown) shared by twins, for the confounding effects are cancelled out when using within-pair differences. In other words, these associations are more likely to be causal, conditional on there being no unmeasured confounders. Except for the three within-pair associations, no association was found for the other seven combinations of risk-predicting measures and risk factors in the within-pair analyses. The null results do not necessarily imply that the observed associations for these combinations are due to familial confounding. Importantly, for all the seven combinations, the confidence interval of the within-pair association in Table 2 contained the point estimate of the association in Table 1. These results suggest that the associations reported in Tables 1 and 2 tend to be alike, and the lack of statistical significance in the within-pair analyses is probably due to the reduced statistical power of using data from twin pairs only.

Given our findings, we hypothesized that DNA methylation-based risk-predicting measures interplay with their associated risk factors in modifying breast cancer risk. Breast cancer studies with data of both risk factors and DNA methylation collected, appropriate design and analytic methods are required to test this hypothesis. Findings from investigating hypothesis might provide novel evidence for the aetiology, early detection and prevention of breast cancer.

The main strengths of our study include a comprehensive assessment of environmental breast cancer risk factors, investigating epigenetic age acceleration using three DNAm age estimators, and the use of within-pair analysis to investigate the role of familial confounding underlying the observed associations. One limitation is that no multiple testing adjustment was performed, and our results should therefore be interpreted with this in mind.

In conclusion, in our exploratory study we found several lifestyle and hormone-related breast cancer risk factors being associated with DNA methylation-based measures that *could* predict breast cancer risk. The associations of epigenetic age acceleration with BMI, alcohol drinking and age at first live birth are unlikely to be due to familial confounding. The interplay between the associated risk factors and risk-predicting measures in modifying disease risk require further investigation.

## Acknowledgements

We thank all women participating in this study. This research is facilitated through access to Twins Research Australia, a national resource supported by a Centre of Research Excellence Grant (ID: 1079102) from the National Health and Medical Research Council (NHMRC). The Australian Mammographic Density Twins and Sisters Study is supported by NHMRC (grant numbers 1050561 and 1079102), Cancer Australia and National Breast Cancer Foundation (grant number 509307). MCS is a NHMRC Senior Research Fellow. JLH is a NHMRC Senior Principal Research Fellow. SL is a Cancer Council Victoria Postdoctoral Research Fellow.

## Compliance with Ethical Standards

This study is approved by Human Research Ethics Committee at the University of Melbourne. All participants have given written informed consent.

## Conflict of interest

The authors declare that they have no conflict of interest.

## Data availability statement

The DNA methylation dataset analysed is available on Gene Expression Omnibus (GEO) under the accession number GSE100227.

